# Dual-Specific Antibody Design Using Artificial Intelligence

**DOI:** 10.64898/2026.08.03.742397

**Authors:** Michael Peer, Inbar Amit, Yael Diesendruck, Ziv Erlich, Yonit Ben David, Meital Gadrich, Nino Oren, Tzvika Hartman, Sharon Fischman, Guy Nimrod, Marek Strajbl, Avi Haleva, Reshef Shilon, Yehezkel Sasson, Reut Barak-Fuchs, Itzhak Meir, Liron Danielpur, Yuval Mor-Scheerer, Nitzan Dubovski, Tal Vana, Dagan Hadar, Anna Voropaev, Yair Fastman, Yanay Ofran

## Abstract

Multibodies, or “two-in-one” Immunoglobulin G (IgG) antibodies, are standard symmetrical IgG molecules engineered to competitively bind more than one antigen within a single variable fragment (Fv) binding surface. This format merges the functional advantages of bispecifics, such as multi-target binding and dynamic adaptation to target concentrations, with the superior manufacturing, developability, pharmacokinetics, and avidity of monospecific IgGs. Moreover, the co-accommodation of multiple paratopes on a single set of 6 CDRs introduces new functional possibilities that can improve efficacy and safety. Multibodies can, therefore, be thought of as force multipliers: for any format of antibodies, or fragments thereof, multibodies can bind double the number of epitopes compared to standard antibodies. While these advantages were recognized more than 15 years ago, the systematic design of multibodies has been intractable due to the challenge of optimizing two binding specificities into one Fv region, without having one of them compromising the other and without inducing poly-reactivity. To overcome this engineering barrier, we have developed an artificial intelligence (AI)-assisted computational platform that enables the design of functional multibodies against virtually any pair of targets. We applied the platform to design nine multibodies combining 15 different unrelated targets. We obtained therapeutic-grade multibodies that bind each desired pair of targets. We demonstrate that the generated multibodies possess excellent developability, high affinity, and stringent specificity, comparing favorably to clinical monospecific benchmarks. Critically, we show that these multibodies exhibit superior functional activity across a diverse range of mechanisms of action (MOAs), including internalization, T-cell engagement, and immune system modulation. This capability to reliably engineer versatile multibodies opens a new domain in antibody therapeutics, enabling complex multipharmacology and novel functions within a natural, cost-effective, and highly developable format. Two of these multibodies are currently in IND enabling studies, with first in human studies expected in 2026. The timeline from idea to a fully optimized, developable, lead candidate, ready for IND enabling studies, is 9 months.

## Introduction

Monospecific IgG antibodies represent the dominant class of biotherapeutics due to their simple manufacturing, robust developability, and long serum half-life. Bispecific antibodies (BsAbs) offer a functional advantage by targeting two distinct antigens simultaneously, yet most clinical-stage BsAbs utilize complex, unnatural formats (e.g., asymmetric chains, scFv/VHH fusions) that frequently incur significant challenges related to CMC (Chemistry, Manufacturing, and Controls) complexity, challenging formulation, shorter half-life, immunogenicity risk, and suboptimal developability profiles^1–3^. For example, lunsekimig, a multispecific construct that antagonized both TSLP and IL-13 and was introduced by Sanofi, has been reported to generate anti-drug antibodies (ADA) in over 43% of the subjects after three doses^4^. Analysis of half-life in humans indicates that the average half-life of Fc-containing bispecifics is ∼14 days, shorter by a week than the reported ∼21 days average of a symmetrical standard IgG^3,5^.

Multibodies, sometimes referred to as two-in-one antibodies, provide a compelling solution. As illustrated in **Figure 1**, multibodies retain the symmetrical (bivalent), native IgG structure, but their Fv binding surfaces are engineered to recognize and competitively bind more than one distinct epitope, which may lie on different antigens. This architecture allows a single arm to bind either target A or target B, granting the molecule the ability to adapt its binding configuration, with either both arms binding target A, both arms binding target B, or each arm binding a different target, based on the local concentrations of the targets.

**Figure 1:**
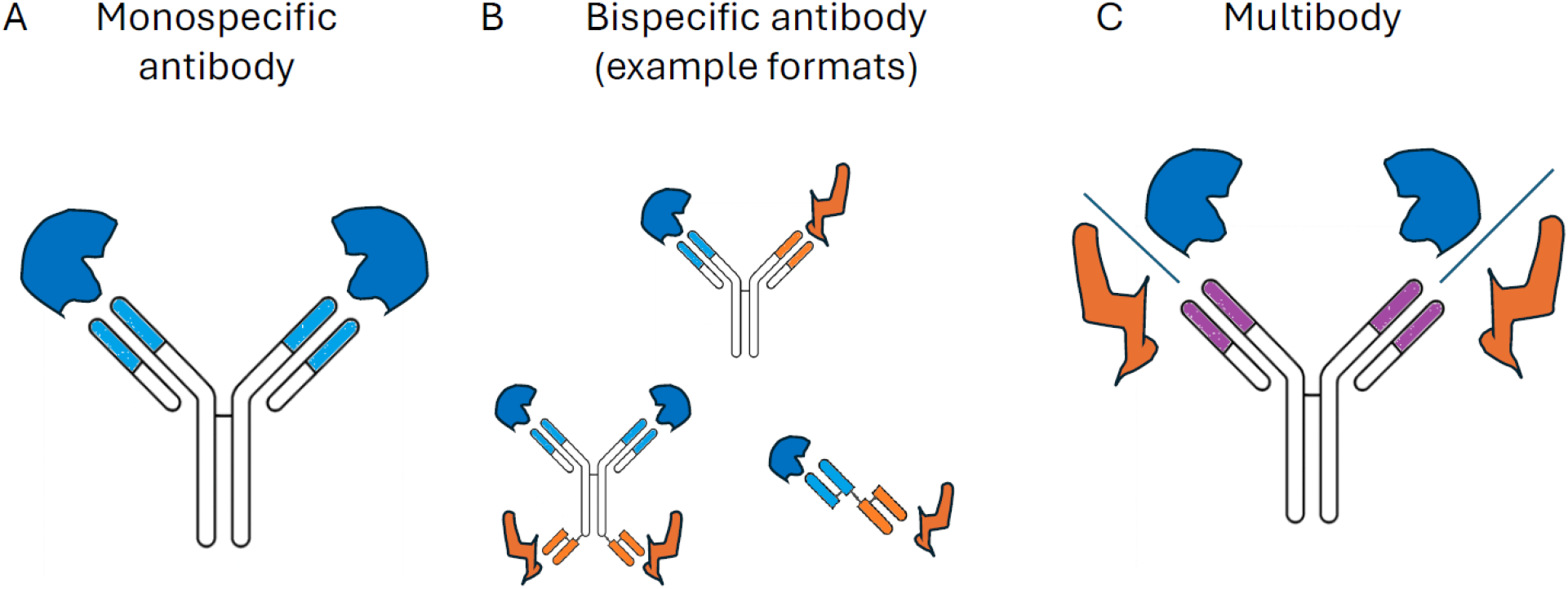
Monospecific antibodies, bispecific antibodies, and multibodies. A) Monospecific IgGs bind only one antigen per arm. B) Bispecific antibodies achieve dual targeting through complex, often unnatural structures. C) Multibodies retain the natural IgG structure while achieving dual specificity within the single Fv of each arm, allowing competitive binding of either target A or B.

Despite their promising attributes, including avidity, dynamic adaptation to target concentrations, and manufacturing simplicity, multibody design has remained largely anecdotal^6–9^ and has not resulted in active clinical candidates. The core challenge lies in simultaneously optimizing the Fv region to accommodate two distinct, potentially overlapping paratopes with high affinity and specificity, all while avoiding non-specific binding (polyreactivity).

Figure 2 illustrates the pharmaceutical, functional, and logistical advantages offered by the multibody format compared to conventional bispecific antibodies. Critically, the symmetrical IgG structure of the multibody is preserved, guaranteeing superior drug-like properties such as high thermal stability, low immunogenicity risk, and, most importantly, a prolonged serum half-life.

From a functional perspective, the multibody is designed for **dynamic adaptation**; each arm competitively binds all its targets. This allows the molecule to adapt its function in tissues or microenvironments driven by the relative local target concentrations, which is vital for targeting heterogeneous expression across tumors and patients. This format also leverages **avidity**—the simultaneous binding of both targets in both arms—to improve apparent affinity, enhance receptor neutralization, and reduce drug sensitivity to target downregulation. Furthermore, this competitive binding enables “XOR gate” functionality, limiting immune overactivation when used in T-cell engagers and opening the way for novel exchange functions.

**Figure 2:**
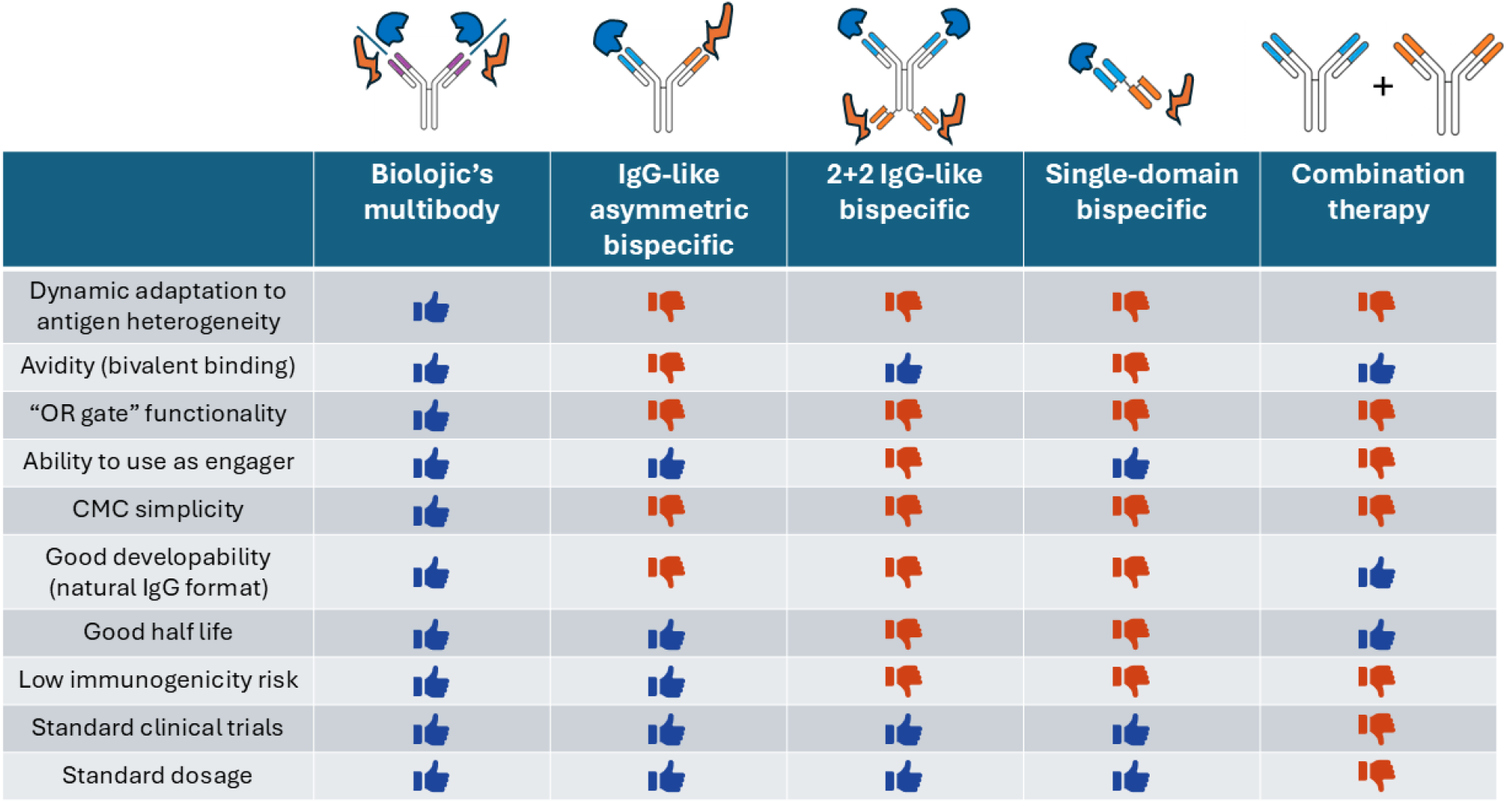
the unique advantages of multibodies compared to other bispecifics. Multibodies enable binding two targets with avidity and dynamic adaptation to target concentrations, while avoiding the complex manufacturing, developability problems, and clinical issues that plague other bispecifics and combination therapies.

The inherent symmetry of the multibody also simplifies purification and large-scale manufacturing processes (simple CMC), making it a significantly more cost-effective and developable therapeutic modality than highly engineered asymmetric bispecific IgG variants. The simplicity of the CMC also reduces timelines and cost of production by ∼50%.

Finally, addressing the growing need for multipharmacology approaches in complex diseases, the multibody format allows for **higher-order combinations**. Combining more than two targets conventionally requires the development of new molecular formats, with the CMC, PK and immunogenicity challenges they may pose, or the combination of separate molecules, complicating cost, timelines, regulatory path and administration. The multibody format enables the combination of two different multibodies into a bispecific format, resulting in a functional tri- or tetra-specific molecule.

## Results

### AI-Enabled Multibody Engineering Platform Overview

Figure 3 shows the multibody engineering platform. It is structured as a four-stage, alternating AI-experimental workflow designed to efficiently navigate the vast sequence space required for dual-specificity. The platform workflow is as follows (see Methods for additional details):

**Figure 3:**
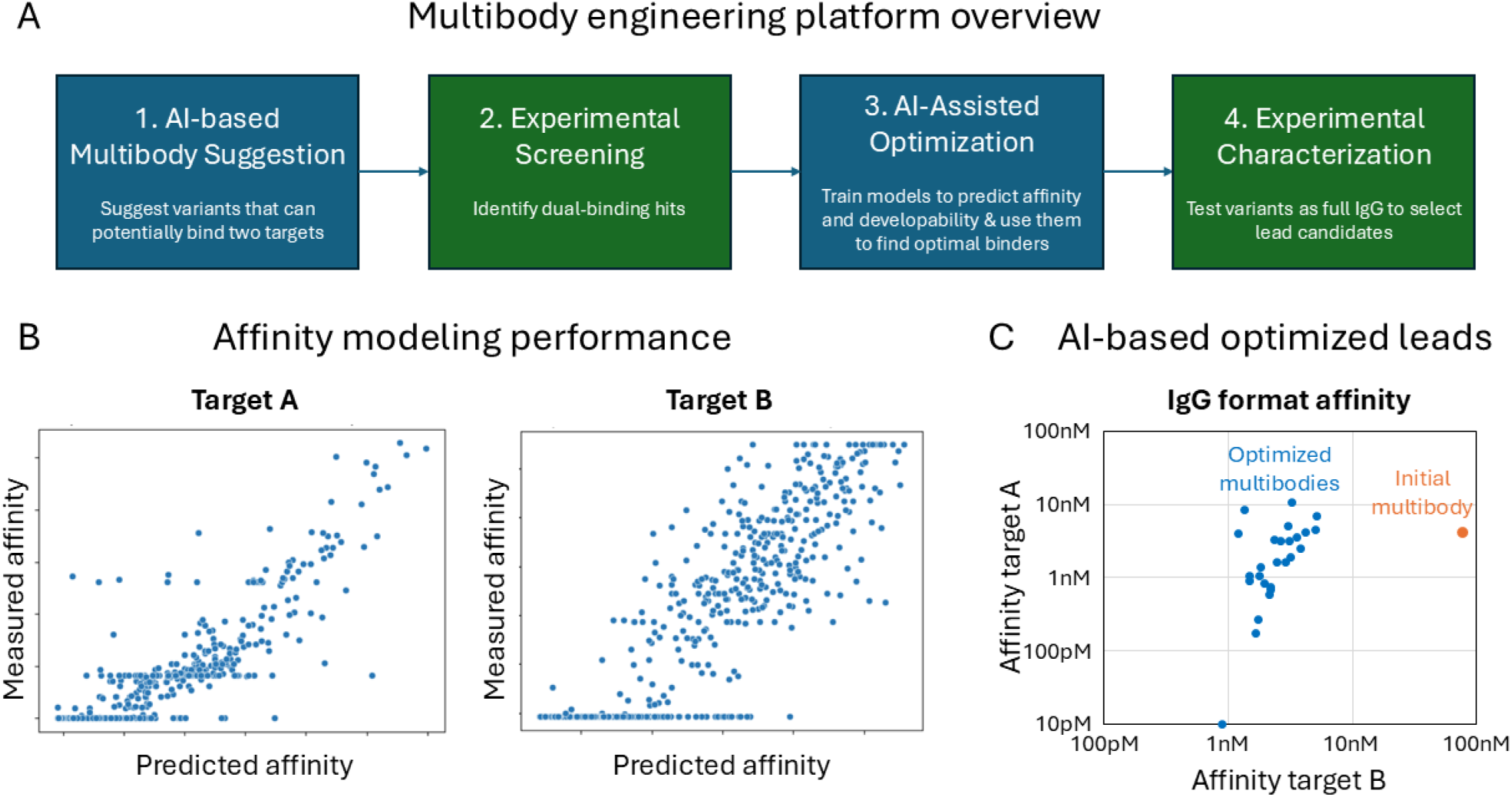
Biolojic Design’s multibody engineering platform. A) To design new multibodies, our platform alternates between AI-assisted design steps (blue) and experimental steps (green): 1. proprietary AI algorithms, tuned by user input, suggest sequences that can potentially bind both targets. 2. the sequences are experimentally screened to identify actual multibodies. 3. multi-objective AI algorithms suggest variants with improved affinity to both targets and good developability. 4. User-selected optimized variants are tested to select final lead IgG candidates. B) Examples of predicted vs. measured affinity of the affinity optimization AI algorithm (step 3), on unseen data. Affinity is measured with proprietary high-throughput evaluation technology. C) Experimental characterization of predicted optimized multibodies (step 4; affinity measured using BLI Kd in IgG format; affinity scale is logarithmic). The pre-optimized multibody from step 2 (orange) had an affinity of 80nM and 4nM to the two targets. Our models suggested multiple optimized multibodies in step 3 (blue). As can be appreciated, optimized multibodies obtained single-digit or sub-nanomolar binding to both targets, with up to two orders of magnitude affinity increase for each target.

1. **AI-based Multibody Suggestion:** Proprietary AI models receive expert user input on targets and desired antibody properties, and propose a diverse library of variants that can potentially bind two pre-specified targets (Figure 3A).
2. **Experimental Screening:** The suggested sequences are experimentally screened to identify initial “hits” – a diverse set of dozens of multibodies that bind both targets, typically with weak to moderate affinity (Figure 3A).
3. **AI-Assisted Multibody Optimization:** A subset of promising hits is selected by the user for multi-objective optimization, and libraries are designed based on these hits. We typically do not expect that the libraries designed at this stage will contain an optimized multibody. Rather, we expect that the libraries will contain a large number of variants that bind one of the targets with high affinity. The libraries are screened separately against each target, to identify diversified binders to each of the targets. Based on the data collected from these screens and on user-selected parameters, for each target separately, we train a model specifically to predict binding affinity of the antibody variants to that target. Similarly, we can collect data on stability, expression and other parameters and train additional models. As shown in Figure 3B, the predicted affinity for both Target A and Target B strongly correlates with experimentally measured values on unseen data. Finally, we search the sequence space for variants that are predicted to have high affinity to both targets and to have a good developability profile.
4. **Experimental Characterization:** Variants predicted to have improved affinity, high yield, and good developability are selected by the user and are then expressed, purified, and tested experimentally to select the final development candidates. An example optimization campaign shown in Figure 3C demonstrates the platform’s performance: a sub-optimal initial dual binder (4nM to target A, 80nM to target B) was optimized to yield multiple variants with dramatically improved affinity. All optimized variants showed comparable or up to 100-fold improved affinity to both targets, with affinities in the single-digit nM or sub-nM range.

### Generation of High-Affinity Multibodies for Diverse Targets

To validate the platform’s reliability and versatility, we executed nine independent multibody engineering campaigns against diverse target pairs (Table 1). The target set included challenging targets such as G-protein coupled receptors (GPCRs), transmembrane proteins, and small cytokines. Furthermore, campaigns were initiated with pre-defined mechanisms of action (MOAs) and epitope requirements. For example, campaigns #1 and #2 targeted the same target pair (Trop-2 and Nectin-4) but were optimized for different epitopes with different functional endpoints: tumor cell internalization (antibody-drug conjugate candidate, #1) versus T-cell engagement/immune synapse formation (T-cell engager tumor arm, #2). Some of these campaigns used earlier versions of our computational development pipeline and some used the current pipeline as described in Figure 3.

**Table 1.** Multibody engineering campaign results. Representative results from a diverse set of campaigns targeting different antigens and optimizing for specific functional mechanisms. Affinities were measured in IgG format using ELISA-EC_50_ (multibodies #2, #4-6, #8-9), SPR (multibodies #1, #7), or flow cytometry EC_50_ (multibody #3; see Methods section for assay details).

| Campaign / multibody | Target A | Target B | Intended MOA | Affinity target A | Affinity target B |
| --- | --- | --- | --- | --- | --- |
| 1 | Trop-2 | Nectin-4 | Tumor cell internalization | 3.1 nM | 17.9 nM |
| 2 | Trop-2 | Nectin-4 | T cell engagement (tumor arm) | 0.4 nM | 1.4 nM |
| 3 | TNFR superfamily target | GPCR superfamily target | T cell engagement (tumor arm) | 7 nM | 21 nM |
| 4 | Immune cell target | Immune cell target | Immune cell engagement (immune arm) | 3 nM | 0.4 nM |
| 5 | PD-1 | GITR | Immune system agonism | 16 nM | 4 nM |
| 6 | PD-1 | OX40 | Immune system agonism | 0.6 nM | 24 nM |
| 7 | TSLP | IL-13 | Immune system antagonism | 0.003 nM | 0.01 nM |
| 8 | Undisclosed target 1 | Undisclosed target 2 | Undisclosed | 0.1 nM | 2.3 nM |
| 9 | Undisclosed target 3 | Undisclosed target 4 | Undisclosed | 0.2 nM | 2.1 nM |

All nine campaigns were successful, obtaining multiple multibodies that met the desired, predefined affinity thresholds for both targets and the developability criteria.

We next benchmarked the binding affinity of our multibodies against relevant clinical-stage monospecific antibodies (comparative ELISA-EC_50_ assays, Figure 4). In all nine cases, the dual specificity of the multibodies did not come at the expense of binding strength to each individual target. The multibodies achieved binding affinities to each individual target that were comparable to, and in some cases superior to, the clinical benchmark antibodies targeting the same epitope. This confirms that the AI-assisted design platform successfully enabled the integration of high-affinity binding sites to both targets in the same antibody.

**Figure 4:**
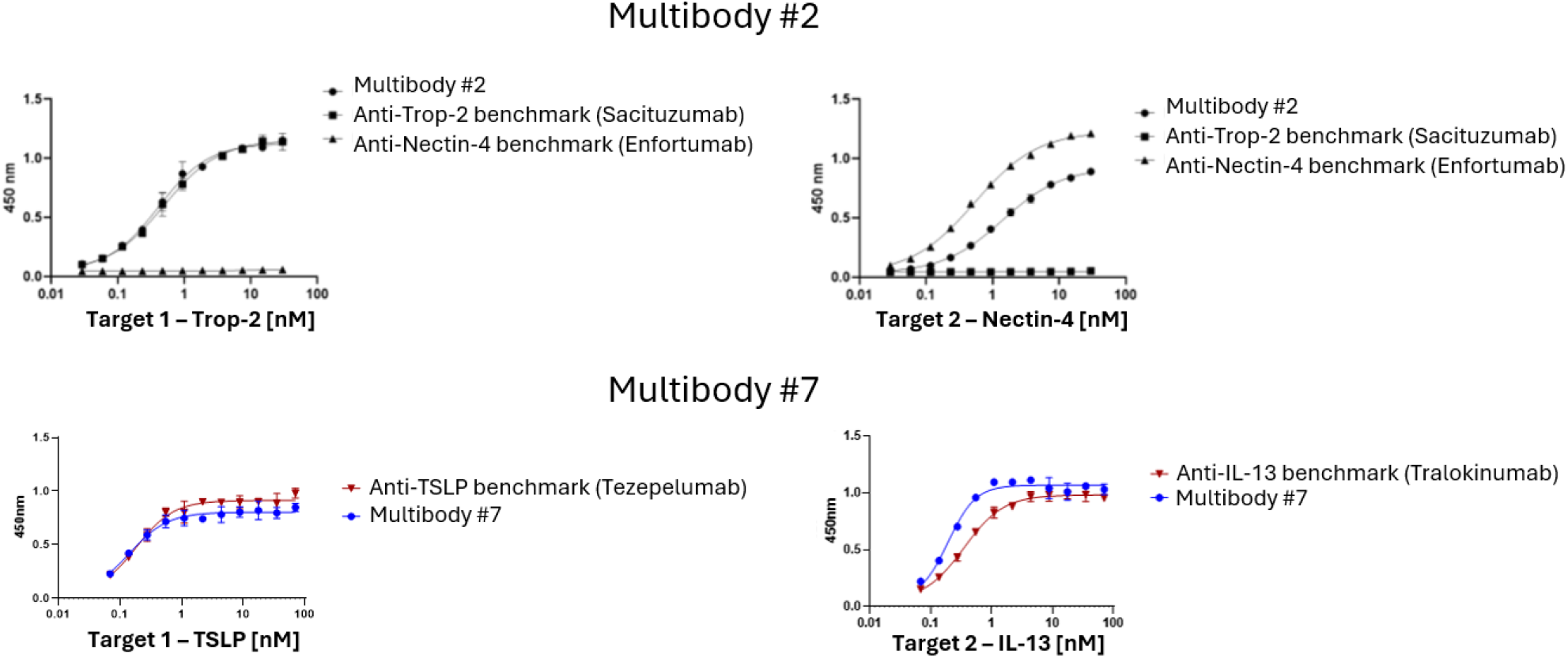
Affinity of multibodies compared to approved monospecific clinical benchmarks. ELISA-EC_50_ binding curves for representative multibodies (#2, #7) demonstrate that their affinity toward each target is comparable to corresponding clinical-stage mono-specific antibodies.

### Developability and Specificity Profile

A critical concern for any engineered therapeutic is its biophysical stability and specificity. Figure 5 shows an assessment of key developability indicators of multibodies: specificity/polyreactivity, hydrophobic interactions, and thermostability. Developability characterization of AI-suggested leads from 4 multibody campaigns (200 multibodies overall, experimentally tested in IgG format) demonstrates that most of them have good specificity, hydrophobicity and thermostability profiles (Figure 5A). Figure 5B shows the results of an example multibody (Multibody #1) tested in the same experiment, with three assays: 1) Baculovirus Particle assay (BVP), used to assess polyreactivity (non-specific binding); 2) Hydrophobic Interaction Chromatography (HIC), used to assess hydrophobic interactions; and 3) Differential Scanning Fluorimetry (DSF), used to assess thermal stability. The results demonstrate that multibody #1 possesses specificity, hydrophobicity and thermal stability within acceptable clinical ranges, and better than several other marketed therapeutic antibodies. These results confirm that multibodies maintain the developability profile characteristic of clinical IgGs, contrasting with the challenges often observed in complex bispecific formats.

**Figure 5:**
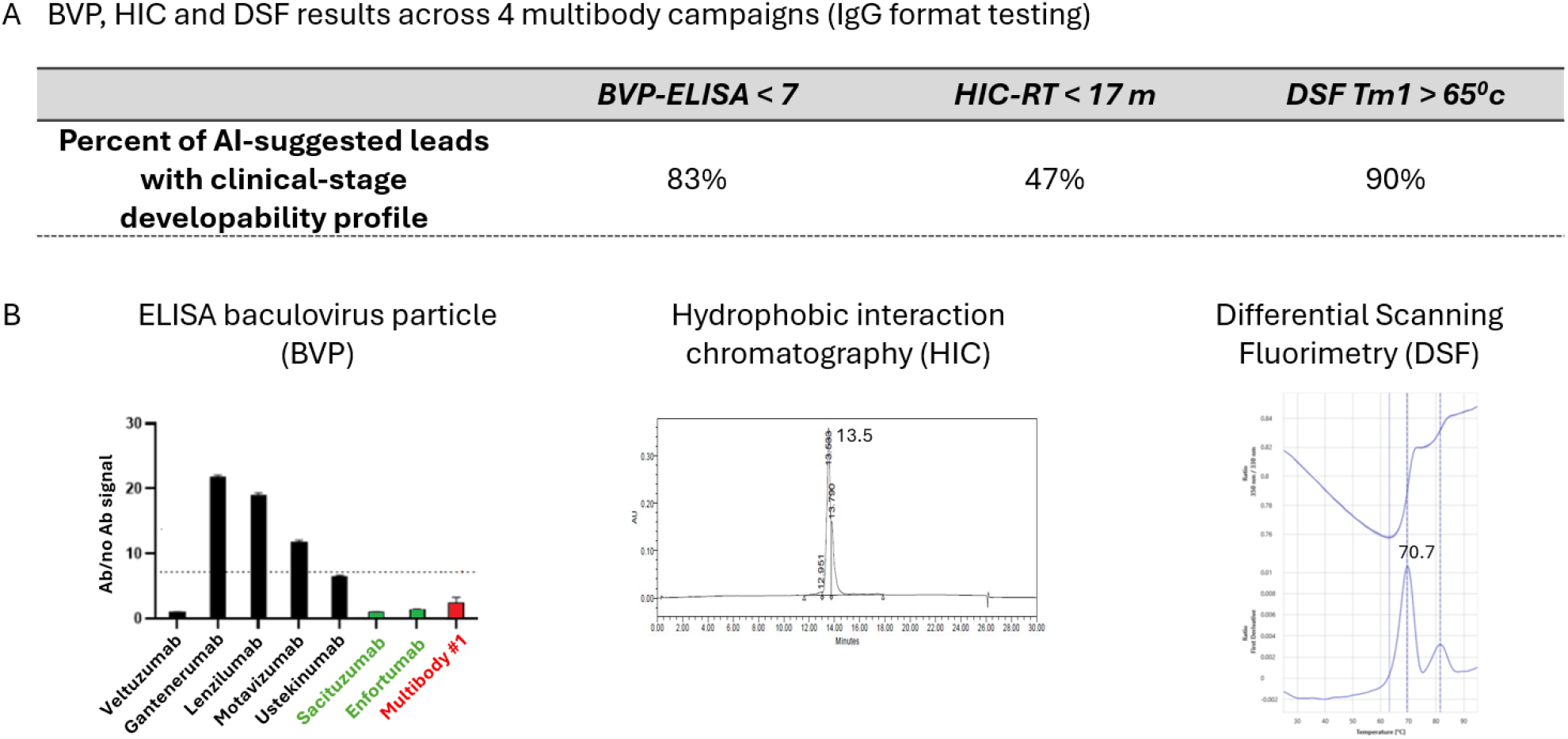
The engineered multibodies possess an excellent developability profile. A) Results of developability testing across AI-suggested leads from 4 multibody campaigns (total of 200 multibodies, tested in IgG format), demonstrating that most leads have a good developability profile including specificity (BVP-ELISA scores), hydrophobicity (HIC retention time) and thermal stability (DSF Tm1). B) Example developability profile of multibody #1. Baculovirus Particle (BVP) ELISA demonstrates specificity (red) similar to clinical benchmarks (green) and better than several clinical therapeutics (black). HIC chromatograms and DSF plots demonstrate acceptable retention times and thermal stability. See Methods section for assay details.

### Superior Functional Activity Across Therapeutic Modalities

We hypothesized that the combined properties of dual competitive binding, avidity, and dynamic adaptation would confer superior functional activity to multibodies compared to monospecific or combination therapies. Each of the multibodies in this PoC study were designed to act through a specific MOA that required a specific set of molecular characteristics. Specifically, some were designed as multibody drug conjugates, with the goal of outperforming mono-specific ADCs. Others were designed to be super engagers, with the goal of outperforming T cell engagers (TCEs) that target a single tumor associated antigen (TAA). Other multibodies were designed to antagonize soluble cytokines, optimizing dose and affinity to outperform monospecific anti-cytokine antibodies. We tested this across three key therapeutic MOAs. The results of these analyses are presented in Figure 6.

**Figure 6:**
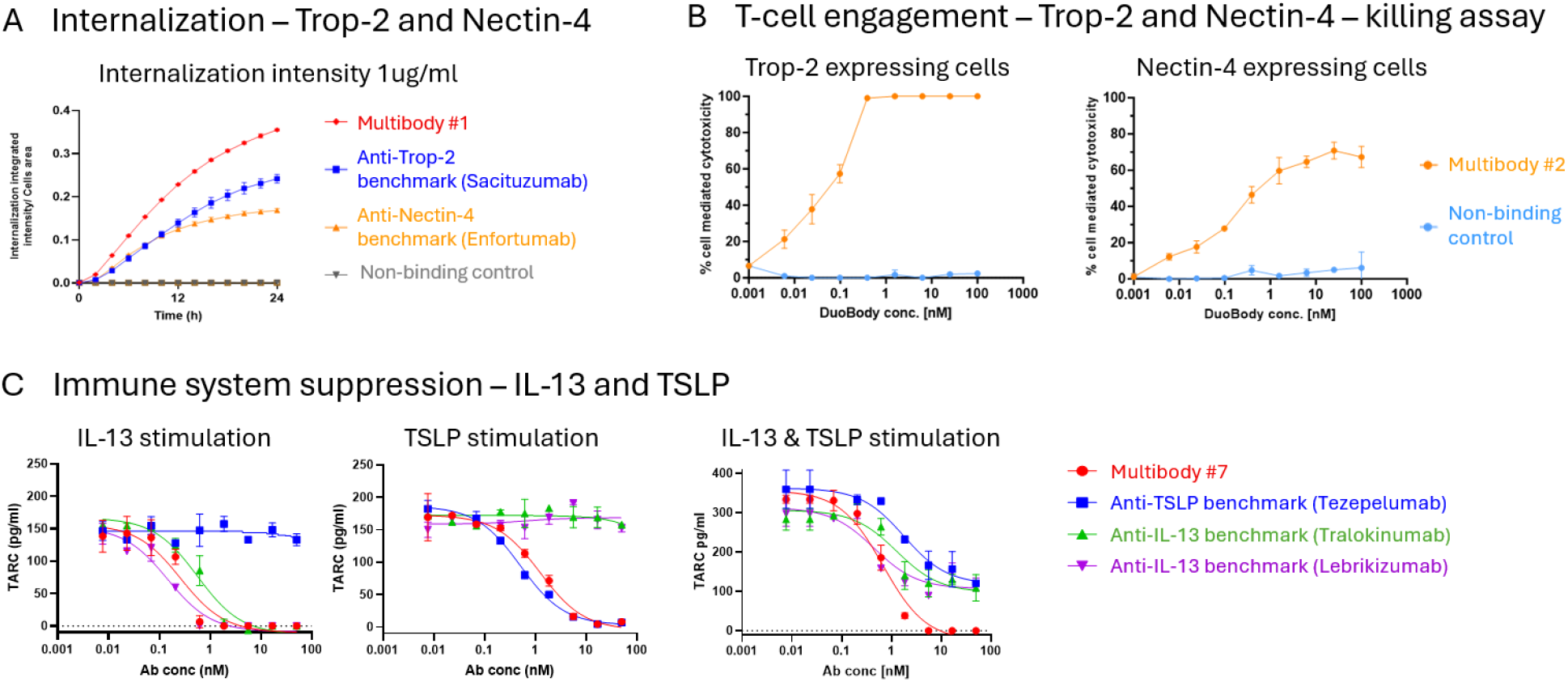
Multibodies surpass approved therapeutic benchmarks across a wide range of functions. A) Multibody #1 demonstrates increased integrated internalization intensity compared to monospecific benchmarks against the same targets. B) A Multibody-based T-cell engager (Duobody format with CD3 binder) induces cytotoxicity in cells expressing either Trop2 or Nectin4, demonstrating target adaptability. C) Multibody #7 maintains complete immune suppression when stimulated by both IL-13 and TSLP, outperforming monospecific clinical benchmarks. See Methods section for assay details.

#### 1. Antibody-Drug Conjugate (ADC) Internalization (Multibody #1)

Multibody #1 (anti-Trop-2, anti-Nectin-4) was designed to target epitopes conductive to internalization. In cells co-expressing both targets, this multibody demonstrated significantly higher internalization rates than the corresponding monospecific clinical benchmarks (Sacituzumab and Enfortumab), suggesting improved delivery of conjugated cytotoxic payload (Figure 6A). Importantly, it has been shown that effective internalization often requires that the

ADC crosslinks two subunits of the receptors. This, obviously, can only be achieved by a bivalent antibody. Hence, while a traditional BsAb that is used as an ADC may internalize well into cells that express both TAAs, it will perform less well on cells where one of the TAAs is downregulated. A multibody-drug conjugate, on the other hand, will retain bivalent binding, and the resulting crosslinking, on all cells that still express one of the TAAs. This enhancement is attributed to the bivalent/avidity effect on the single-chain variable fragment (scFv) binding site.

#### 2. T-Cell Engagement (Multibody #2)

Multibody #2 (anti-Trop-2, anti-Nectin-4) was engineered as the tumor-targeting arm of a Duobody-formatted T-cell engager (TCE) by fusing it to an anti-CD3 binder. This required binding different epitopes on each of the TAAs compared to multibody #1, which was optimized for internalization instead of T-cell engagement. When tested against two cell lines—one expressing Trop-2 and one expressing Nectin-4—the multibody-based TCE induced potent, dose-dependent, T-cell-mediated cytotoxicity in both cell lines (Figure 6B). This demonstrates the multibody’s capacity for dynamic adaptation, allowing it to effectively engage T-cells regardless of which target is prevalent on the tumor cell surface, a significant advantage over a bsAb that might be limited if its TAA target is absent. Note that in this case it is not straightforward to replace the multibody with a bsAb, as this will require a new, untested format with at least 3 Fvs.

#### 3. Immune System Modulation (Multibody #7)

Multibody #7 was designed as an antagonist against two proinflammatory cytokines, IL-13 and TSLP, relevant in inflammatory diseases. When only one cytokine was present, assays measuring the suppression of TARC secretion in response to IL-13 or TSLP stimulation showed that the multibody was comparable to the monospecific benchmarks (Tezepelumab and Tralokinumab / Lebrikizumab; Figure 6C, left and center panels). Crucially, when immune cells were stimulated with both IL-13 and TSLP, as is the case in patients, Multibody #7 maintained complete immune suppression, while the monospecific benchmarks showed only partial suppression (Figure 6C, right panel). This result highlights the ability of the multibody to serve as a stronger, more comprehensive immune modulator than existing state-of-the-art combination strategies.

## Discussion

The computational platform described herein provides a robust, generalizable solution to the long-standing challenge of engineering dual-specific antibodies in the natural IgG format. Our results demonstrate that the AI-guided workflow reliably generates development candidates with a 9/9 success rate across nine diverse campaigns, targeting a range of medically relevant, and often difficult, targets including GPCRs and small cytokines.

The key finding is that these multibodies successfully decouple the structural complexity of dual-targeting from the resulting drug-like properties. By retaining the standard IgG structure, our candidates exhibit impeccable developability (low BVP score, acceptable HIC, high thermal stability) and pharmacokinetics, resolving the key liabilities associated with many non-IgG bispecific formats. Furthermore, the capacity for competitive binding within a single Fv arm confers significant functional advantages. The observed superior internalization rates, dynamic tumor cell recognition by the T-cell engager, and complete immune suppression in the presence of two stimuli collectively demonstrate the enhanced clinical utility of this format over current state-of-the-art mono- and bispecific therapeutics.

The capability to rapidly and reliably engineer these molecules opens a new domain of precision medicine. The dynamic, concentration-driven binding profile of multibodies allows them to adapt their function to heterogeneous antigen expression across tissues, tumors, and patient populations, offering a route to improved efficacy and reduced dose-related toxicity compared to fixed-ratio combination therapies. Two of the multibody candidates described are currently advancing toward IND, with projected human trials commencing in 2026. This technical achievement establishes the multibody as a promising next-generation therapeutic format poised to disrupt existing clinical practice across numerous indications.

## Methods

### AI-enabled multibody engineering platform

The multibody engineering platform receives as input the targets’ sequences and/or structures, along with expert user input on targets’ features and desired multibody properties. The platform then follows a four-stage workflow to obtain functional multibodies that can each bind both targets. The workflow is described below, although full AI model architecture and parameters are not disclosed due to their proprietary nature:

1. **AI-based Multibody Suggestion:** user-defined inputs (desired targets’ sequences and/or structures and user-selected parameters) are fed to a multibody generation model. This generative model consists of a protein large language model (LLM) component and a structural modeling component, used together to represent the antibody and target structure and generate multibody sequences that are predicted to bind both targets.
2. **Experimental Screening:** The suggested multibody sequences are expressed as a library and experimentally screened to identify hits – multibodies that bind both targets.
3. **AI-Assisted Multibody Optimization:** the optimization consists of three sub-stages:
  ∘ **Data generation** – a subset of multibody sequences is selected from the previous stage, and libraries are designed by human experts through selection of mutations to these sequences. These libraries are expressed and displayed to collect data on each sequence’s affinity to each target, as well as stability, expression and other parameters of interest.
  ∘ **Model training** – based on the labelled data collected from these screens and on user-selected parameters, models are trained to predict the binding affinity of the antibody variants to each target, as well as their stability, expression and other developability parameters. Each of these models consists of an encoding layer that represents antibody properties and a prediction head that predicts the property of interest (affinity to each target and various developability parameters).
  ∘ **Sequence search** – the trained models are used in a multi-objective search of the vast potential sequence space. The goal of this search is to identify variants that satisfy all desired constraints simultaneously – high affinity to both targets and good developability profile across multiple developability parameters.
4. **Experimental Characterization of Selected Leads:** the user selects variants that are predicted to have improved affinity and developability. These variants are expressed and tested to select the final development candidates.

### Affinity measurement

affinity was tested in IgGs, except for multibodies #5-6 that were tested in yeast display.

- **ELISA-EC**_**50**_ was performed by coating antigen A or antigen B on a plate and titrating the antibody in a range of concentrations. Detection was performed using anti-His-HRP antibody and absorbance was measured at 450nm.
- **SPR analysis** was performed using Biacore T200 and S200 systems. A CM5 sensor chip was immobilized with anti-human Fc antibody to capture the tested multibody. Antigen injections were performed in a multi-cycle kinetics format at a range of concentrations. Kinetics were calculated using 1:1 binding.
- **Flow-cytometry EC**_**50**_: monoclonal antibody (Ab) EC_50_ values were determined by incubating target-expressing cells with serial dilutions of the tested Abs. Following incubation, cells were washed twice and stained with anti-human Fc-647 detection Ab. Binding was measured using a CytoFLEX flow cytometer, and EC_50_ values were calculated by fitting the data to a two-parameter model in GraphPad Prism.

### Developability assays

- **BVP analysis** was performed by coating high-binding 96-well ELISA plates with BVP and applying 1000 nM of tested antibodies. Detection was performed using anti-Fc-HRP antibody and absorbance was measured at 450nm.
- **HIC analysis** was performed by spiking 0.6mg of tested antibody into mobile phase A solution (2M ammonium sulfate and 0.1M sodium phosphate at pH 6.5) in 1:1 v:v ratio before the analysis. Antibody was injected to a Proteomix HIC Butyl-NP5 5um Non-Porous 4.6*35mm column (Sepax) at a volume of 40ul (a total of 12 ug), and elution was performed in a linear gradient of mobile phase A and mobile phase B (0.1M sodium phosphate at pH 6.5) over 20 min from 90% solution A to 100% solution B at a flow rate of 1 ml/min. Results were analyzed using 280nm.
- **DSF analysis** was performed using NanoDSF Prometheus Panta (NanoTemper) at the Weizmann institute. The antibody was diluted to a concentration of 0.5 mg/mL in PBS pH 7.4 loaded on capillaries (PR-C002, NanoTemper) and measured in duplicates using excitation power of 93%, with temperature gradient range of 25°C–95°C at 1°C/min rate. Analysis was performed using PR Panta Analysis V1.4.4.

### Functional assays

- **Internalization:** an in vitro internalization assay was conducted using a cell line co-expressing both target antigens to assess internalization of Multibody #1. A pH-sensitive fluorescent probe was used to detect antibody internalization, as its signal increases in acidic endosomal and lysosomal compartments, allowing visualization of productive trafficking critical for ADC mechanism of action. Internalization kinetics of test antibodies were monitored using the Incucyte® live-cell imaging system. Cells were seeded and treated with test antibodies (1 µg/mL) pre-mixed at a 1:3 molar ratio with Incucyte® Human Fabfluor-pH Antibody-Orange reagent, then incubated at 37°C for 24 h. Images were captured hourly to quantify fluorescence increase associated with antibody internalization into acidic compartments.
- **T-cell engagement (killing assay):** Peripheral blood mononuclear cells (PBMCs) were co-cultured with either HEK293-TROP2 or HEK293-NECTIN4 expressing cell lines for 48h. To determine the level of cytotoxicity, the Cytotoxicity Detection Kit (LDH) (Roche) was used. Cytotoxicity percentage was defined as LDH secreted levels of treated samples relative to maximal LDH secretion.
- **Immune system downregulation (TARC assay):** TARC (Thymus and Activation Regulated Chemokine, CCL17) is a chemokine involved in recruiting Th2 cells and CLA+ CD4+ T cells in inflammatory conditions, that serves as a clinical biomarker for disease severity and treatment efficacy. Both IL-13 and TSLP directly upregulate TARC production by dendritic cells. To measure TARC production, hPBMCs were stimulated for 48h with either IL-13, TSLP or with the combination of both cytokines. Immediately following stimulation, cells were treated with Multibody #7 or clinical benchmarks against the two antigens. TARC levels in the supernatant were measured by ELISA.

